# Single molecule long-read real-time amplicon-based sequencing of *CYP2D6*: a proof-of-concept with hybrid haplotypes

**DOI:** 10.1101/2022.08.17.503990

**Authors:** Rachael Dong, Megana Thamilselvan, Jerome Clifford Foo, Jacqueline DeJarnette, Xiuying Hu, Beatriz Carvalho Henriques, Yabing Wang, Keanna Wallace, Sudhakar Sivapalan, Avery Buchner, Vasyl Yavorskyy, Kristina Martens, Wolfgang Maier, Neven Henigsberg, Joanna Hauser, Annamaria Cattaneo, Ole Mors, Marcella Rietschel, Gerald Pfeffer, Katherine J. Aitchison

## Abstract

CYP2D6 is a widely expressed human xenobiotic metabolizing enzyme, best known for its role in the hepatic phase I cytochrome P450 enzyme system, where it metabolizes ∼20% of medications. It is also expressed in other organs including the brain, where its potential role in physiology and mental health traits and disorders is under further investigation. Owing to the presence of homologous pseudogenes in the *CYP2D* locus and transposable repeat elements in the intergenic regions, the gene encoding the CYP2D6 enzyme, *CYP2D6*, is one of the most hypervariable known human genes - with more than 165 core haplotypes. Haplotypes include structural variants, with a subtype of these known as hybrid haplotypes or fusion genes comprising part of *CYP2D6* and part of its adjacent pseudogene, *CYP2D7*. The fusion genes are particularly challenging to identify. High fidelity (HiFi) single molecule real-time (SMRT) long-read sequencing can cover whole *CYP2D6* haplotypes in a single continuous sequence, and is therefore ideal for structural variant detection. In addition, it is highly accurate and suitable for novel haplotype identification, which is necessary as new *CYP2D6* haplotypes are continuously being discovered, and many more likely remain to be identified in relatively understudied populations such as Indigenous Peoples. The aim of the present work was to develop an efficient and accurate HiFi SMRT amplicon-based method capable of detecting the full range of *CYP2D6* haplotypes including fusion genes. We report proof-of-concept for 24 amplicons including three positive controls, aligned to fusion gene haplotypes, with prior cross-validation data. Amplicons with *CYP2D7-D6* fusion genes, including positive controls, aligned to the *\*13* subhaplotypes predicted (*\*13F*, *\*13A2*) with 100% accuracy, with the exception of one that aligned at 99.9%. Alignment of the *\*68* was 100% and above 99.9% to the *CYP2D6*68* partial sequences EU5300606 and JF307779, respectively. The best alignments for the remaining *CYP2D6-2D7* fusion genes were ≥99.7% (to 3 significant figures). Lower percentage alignment for *CYP2D6-2D7* fusion genes may reflect imperfect PCR optimization and/or the possibility that we may have haplotypes not yet in public databases. Further work on these is in progress. Moreover, we have adapted this method for non-hybrid haplotypes. This technique could therefore suffice for the characterization of the full range of *CYP2D6* haplotypes. The method that we have developed could be extended to other complex loci and to other species in a multiplexed high throughput assay.

## Introduction

CYP2D6 is a xenobiotic metabolizing enzyme widely expressed in multiple organs including the liver, intestine, brain, gonads, and thyroid gland^1–3^. While it is best known for its role in the hepatic phase I drug cytochrome P450 enzyme system, where it metabolizes an estimated 20% of medications^4–6^, it also has physiological roles. In the brain, it is found in cerebellar Purkinje cells and cortical pyramidal neurones^3,7^. It is colocalized with the dopamine transporter^8^, and dopamine transporter inhibitors such as cocaine also inhibit CYP2D6^9,10^. Enzyme activity appears to modulate resting brain perfusion, with suggestive involvement in regions associated with alertness or serotonergic function^11^, and a rodent model developed to further explore this^12^. It is involved in steroid biosynthesis (conducting the 21-hydroxylation of progesterone and allopregnanolone^13^, as well as in the synthesis of dopamine from *m*- and *p*-tyramine^14,15^, and of serotonin from 5-methoxyndolethylamines^16^ including 5-methoxytryptamine (a metabolite and precursor of melatonin^17^). The enzyme is induced in alcoholism^18^ and in pregnancy^19^. There has also been suggestive evidence of an association between enzyme status and personality traits^20–22^.

The gene encoding the CYP2D6 enzyme, *CYP2D6*, lies at chromosome 22q13.2 adjacent to two pseudogenes, *CYP2D7* and *CYP2D8*. The three genes are highly homologous^23–25^, and this, together with transposable repeat elements in the intergenic regions^24^, predisposes the locus to the generation of structural variants and novel mutations. Indeed, with more than 160 core haplotypes (or strings of genetic variants) identified to date^26,27^, *CYP2D6* is one of the most hypervariable known human genes. Variants include single nucleotide variants (SNVs), small insertions and/or deletions (indels), and structural variants^28^. Structural variants include gene duplications and multiplications, complete deletions of the entire gene, and recombination events involving *CYP2D7*^28–32^. The recombination events involving *CYP2D7* result in the formation of hybrid or fusion genes^28–31,33–37^ (Figure 1). *CYP2D7-2D6* fusions have a 5’-portion derived from *CYP2D7* and a 3’-portion derived from *CYP2D6*; these hybrids are non-functional^28^. *CYP2D6-2D7* fusions have a 5’-portion derived from *CYP2D6* and a 3’-portion derived from *CYP2D7*.

**Figure 1.**
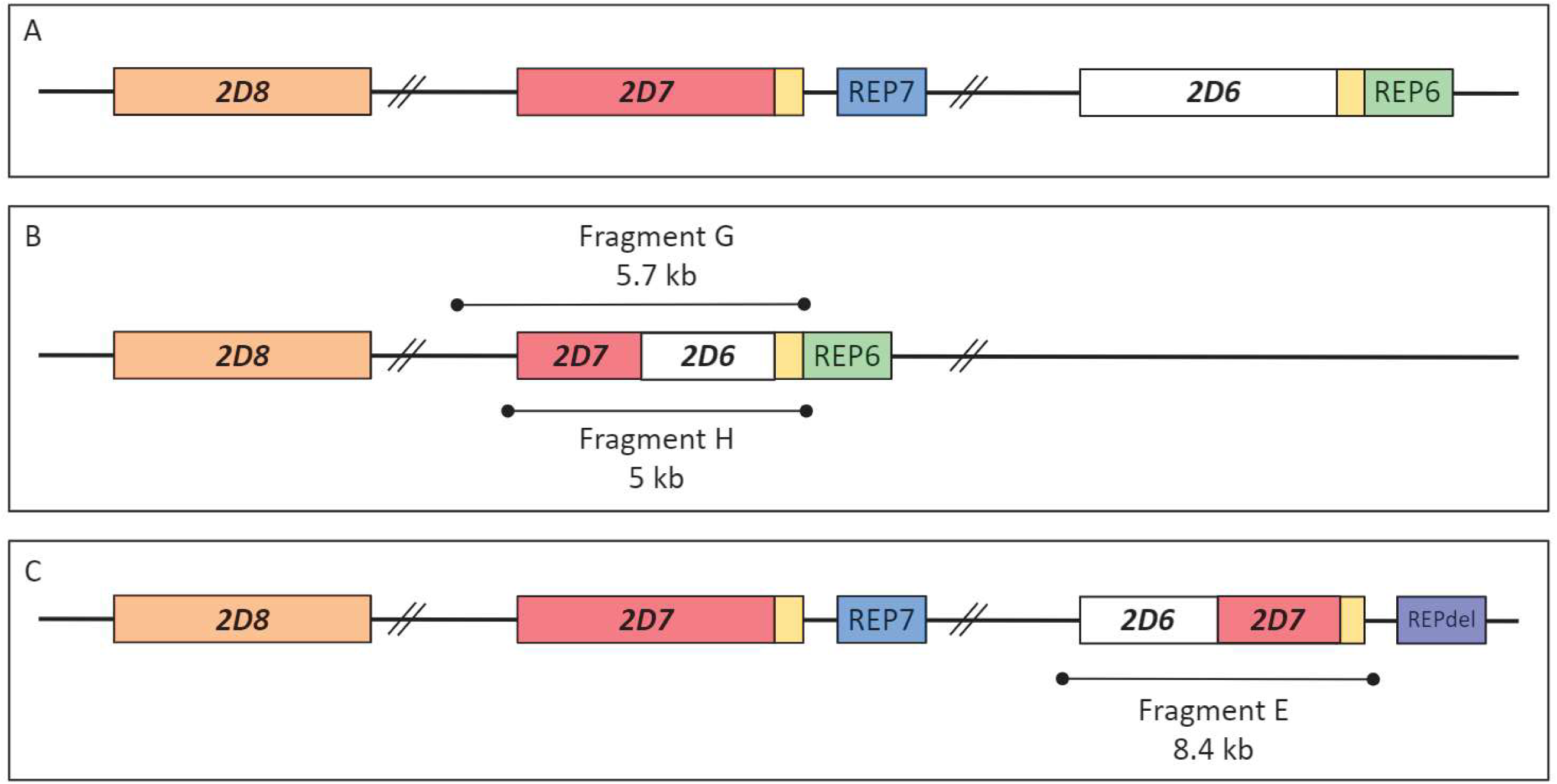
*CYP2D* wild-type locus and fusion genes, with amplicons diagrammed. (**A**) *CYP2D* wild-type locus. (**B**) Example *CYP2D7-2D6* fusion gene. The G and H amplicons are predicted to be 5.7 and 5 kb, respectively. (**C**) Example *CYP2D6-2D7* fusion gene. The E amplicon is estimated as ∼8–8.4 kb.

The Pharmacogene Variation Consortium assigns levels of function (no function, decreased, normal, or increased) to *CYP2D6* haplotypes that correspond to enzyme or phenotype predictions (poor metabolizers, intermediate, normal, or ultrarapid, respectively)^26,38–40^. As an example of haplotype to phenotype prediction, an individual with two no function (or null) haplotypes is a poor metabolizer, with no active enzyme. This has implications for the metabolism of relevant medications. In this example, such individuals are unable to metabolize codeine from its inactive prodrug status to the metabolite with analgesic effect and hence do not experience any analgesic effect with this medication^41^. The *CYP2D6-2D7* fusion or hybrid haplotypes have zero function (e.g., *\*36*, *\*68*, and the ones with the 1847G>A SNV) or uncertain/unknown function(e.g., *\*61* and *\*63*)^26,28^. Hybrid haplotypes are found either as a single haplotype or in tandem with another *CYP2D6* haplotype^28–31,36,37^.

*CYP2D6* hybrid haplotypes are common in the general population, with a frequency estimated as at least 6.7%^42^. In our sample, the Genome-based therapeutic drugs for depression (GENDEP) study^43^, out of 853 patients with depression, the frequency is 2.6% (22/853). Owing to the range of enzyme phenotype corresponding to *CYP2D6* hybrids, their accurate detection is important for predicting prescribing implications^42^, as well as potentially for neuroscience and physiology more generally.

The detection of *CYP2D6* hybrid haplotypes is however, challenging for many genomic technologies, with incorrect and incomplete characterization being commonplace (Carvalho Henriques et al., 2021b). For example, the AmpliChip CYP450 Test did not cover hybrid haplotypes, and hence none of the 19 patients subsequently identified as having hybrid haplotypes had previously had these found, with 2 having been genotyped as *CYP2D6*1/*1* (wild-type) and 4 as ‘no call’^43^. We have previously reported the use of methods including Sanger sequencing to characterize hybrid haplotypes^43–45^. However, Sanger sequencing poses limitations for haplotype phasing (determining which combination of variants lies on which allele) and discriminating whether sequence is derived from *CYP2D6* or *CYP2D7*^46^. As a relatively labour-intensive and time-consuming technique, Sanger sequencing is best suited to low throughput.

Short-read next generation sequencing (NGS) is useful for the detection of SNVs and small indels, but less useful for structural variant detection with haplotype phasing, which requires information across longer sequence or read lengths^47^. Single molecule real-time (SMRT) long-read sequencing provided by Pacific Biosciences (PacBio) and Oxford Nanopore (ONT) can achieve structural variant detection with haplotype phasing. However, until relatively recently, these SMRT technologies had a lower accuracy than short-read NGS. In 2019, circular consensus sequencing (CCS) was optimized to generate highly accurate (99.8%) long high fidelity (HiFi) reads^47^, with a median length of 13.4 kb^48^. At this accuracy level, SNVs and short indels may be identified as well as structural variants. With *CYP2D6* haplotypes being under 10 kb and including all types of variation, this technology is eminently suited for the identification of the full range of haplotypes including novel, unidentified haplotypes. The latter is important as to date, there are populations (e.g., Indigenous peoples) in which this gene is relatively less studied than in others.

Some *CYP2D6* sequencing using HiFi SMRT has already been conducted in previous research. For 25 individuals with prior AmpliChip CYP450 genotype, including four with “*XN*” representing more than one copy of specific haplotypes (e.g., *CYP2D6*1/*2XN*), all genotypes were concordant with the genotype resulting from the SMRT data other than one^49^. In this case, the prior genotype was *CYP2D6*4/*4* and the new genotype was *CYP2D6*4/*5,* with the *CYP2D6*5* representing a complete deletion of the *CYP2D6* gene that had been missed by the AmpliChip (the design of which is now recognized as being able to detect only a subset of *CYP2D6*5* haplotypes^43^). In addition, one novel trinucleotide deletion and one novel SNV were detected in this group of samples and confirmed by Sanger sequencing. SMRT data for *CYP2D6* has also been compared to data from targeted Illumina NGS in 17 individuals^50^. These 17 included one hybrid haplotype (*CYP2D6*36*), including a duplication thereof and its occurrence together with *CYP2D6*10* in a hybrid tandem (*CYP2D6*36+*10*). A recent study has applied the HiFi SMRT technology to 561 patients treated with tamoxifen, and to separate cohorts treated with tamoxifen and venlafaxine^40^. In the tamoxifen-treated dataset, only four individuals with hybrids were identified by the SMRT, and the hybrid haplotype was not specified (Supplementary Table S2^40^). The latest relevant SMRT paper used a long-range polymerase chain reaction (L-PCR) technique that amplified a 6.1 kb region spanning from 712 bp upstream to 1176 bp downstream of the NB_008376.4 *CYP2D6* RefSeq coding sequence, and, in addition (by design), amplified the corresponding region from *CYP2D7*, generating a 7.6 kb amplicon in an analysis of 377 Solomon Islanders^51^. From the SMRT data, 27/365 (7.6%) samples appeared to have a *CYP2D6-2D7* fusion haplotype with breakpoints in exon 8 (consistent with a *CYP2D6*63*), and 7/365 (2%) samples appeared to have *CYP2D7-2D6* fusions (*CYP2D6*13*). However, there was a degree of discrepancy between the above and TaqMan CNV intron 9 and exon 9 data, with not all of the samples with a *CYP2D6-2D7* predicted fusion haplotype having a higher intron 2 than exon 9 CNV count, and 1/7 of the predicted *CYP2D7-2D6* fusions not having a high exon 9 count. In addition, the upstream region covered was insufficient for submission of novel haplotypes to PharmVar, which requires at least 1600 bp upstream of the ATG start sequence (to cover the - 1584C/G SNP).

The research gap that we address herein is therefore: creating an efficient and accurate HiFi SMRT amplicon-based method capable of detecting the full range of *CYP2D6-2D7* and *CYP2D7-2D6* fusion genes, which could be extended to other *CYP2D6* haplotypes and adapted for multigene panels.

## Methods

### Samples

Used herein was the subset of 95 DNA samples from patients with depression in the Genome-based therapeutic drugs for depression (GENDEP) pharmacogenomics clinical trial, specifically, the 19 samples with *CYP2D6* hybrid haplotypes previously described by Carvalho Henriques et al.^43^, plus one additional putative hybrid identified by TaqMan copy number variant (CNV) screening using the methodology described^43^. GENDEP was designed to identify pharmacogenomic predictors of response to two antidepressants, nortriptyline and escitalopram, the metabolism of which both involve CYP2D6^52^. DNA was extracted from venous blood. The 19 samples had prior data from multiple technologies (the AmpliChip CYP450 test, TaqMan SNV and CNV data, the Ion AmpliSeq Pharmacogenomics Panel, PharmacoScan, and Sanger sequencing) resulting in consensus genotype calls, while the one additional sample had only prior AmpliChip CYP450 (*CYP2D6*1/*2*) and TaqMan CNV data that were unequal across different regions of *CYP2D6* (intron 2, intron 6, and exon 9 calls of 3, 3, and 2, respectively), indicating the presence of a *CYP2D6-2D7* fusion gene. In total, there were 20 samples, two of which had consensus genotypes consistent with two hybrid haplotypes. In addition, we used two samples from the Genetic Testing Reference Material Program (GeT-RM) collection as positive controls for *CYP2D6-2D7* (NA18545, *CYP2D6*5/*36x2+*10x2*) and *CYP2D7-2D6* (NA19785*, CYP2D6*1/*13+*2)* hybrid haplotypes, respectively^53^.

### L-PCRs using barcoded universal primers for multiplexing amplicons

We adapted protocols for amplifying hybrid-specific amplicons E, G, and H as described^29–31,43^ for the PacBio barcoded universal primers for multiplexing amplicons method^54^. The E amplicon is for *CYP2D6-2D7* fusion genes, while the G and H cover the *CYP2D7-2D6* fusion genes. A 5’ universal sequence and a 5’ amino modifier C6 (5AmMC6, to prevent unbarcoded amplicons from being sequenced) were added to each primer (Supplementary Table S1, Thermo Fisher Scientific) for the first-round L-PCR.

First-round L-PCRs were performed using KAPA HiFi HotStart ReadyMix (Roche Molecular Systems). Reactions (50 µl, in duplicate) contained primers at 0.3 µM each, dNTPs at 0.6 µM, MgCl_2_ at 2.75 µM, 2% DMSO, and 3 µl of template at 25-202 ng/ µl (more template was used for samples previously showing a relatively low amplicon generating efficiency). For the E amplicon, DNA was amplified for 30 cycles with denaturation at 98°C for 30 s, cycling at 98°C for 10 s, annealing at 85°C for 30 s, and extension at 73°C for 7.5 min (latterly adjusted to 8 min), followed by a terminal extension step of 73°C for 10 min. For the G amplicon, conditions were: denaturation at 98°C for 30 s; 30 cycles of 98°C for 10 s, and annealing and extension at 72°C for 6 min; followed by a terminal extension step of 72°C for 7 min. For the H amplicon, conditions were: denaturation at 95°C for 30 s; 35 cycles of 98°C for 20 s, annealing at 74°C for 30 s and extension at 73°C for 5 min; followed by a terminal extension step of 72°C for 10 min.

PCR products were visualized on a 1% agarose gel, quantified using the Qubit dsDNA HS assay using a Qubit 3.0 Fluorometer (Thermo Fisher Scientific), and visualized using an Agilent DNA 12000 kit on an Agilent 2100 Bioanalyzer (Agilent Technologies) (Figure 2). As some nonspecific products and primer oligomers were seen on the Agilent output for amplicons E and H, respectively, size selection using AMPure PB Beads (PacBio) was used. Purified sample and peak concentrations were measured with the Qubit dsDNA HS and Agilent DNA 12000 assays, respectively.

**Figure 2.**
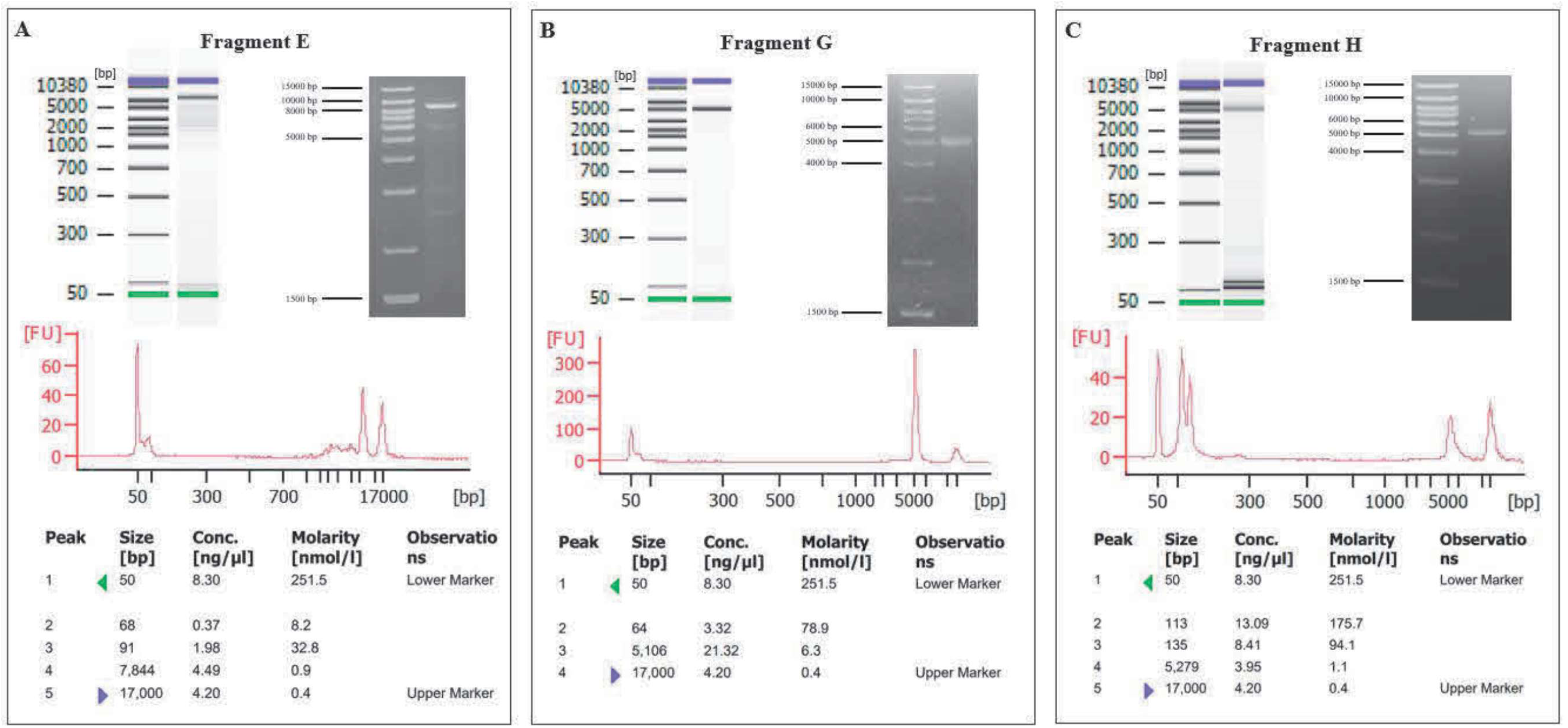
**Quality control of first-round L-PCR by agarose gel and Agilent 2100 Bioanalyzer (DNA 12000 assay) electrophoresis**. (**A**) Representative E amplicon. (**B**) Representative G amplicon. (**C**) Amplicon H. For the Agilent output, the x-axis indicates length in base pairs (bp) and the y-axis fluorescence intensity in fluorescence units [FU]. The lower marker (50 bp) and upper marker (17000 bp) are the first and last peaks, respectively. Electropherogram plots are transformed into automated gel electrophoresis images (top left), where the bottom marker (green), top marker (purple), and PCR products are visualized alongside the ladder (bp). The 1% agarose gel with a 1 kb Plus DNA Ladder (Thermo Fisher Scientific) is shown in the top right.

In the second-round L-PCR, barcoded universal primers (barcoded universal F/R primers plate 96 v2, PacBio) were attached to the forward and reverse universal sequences in the amplicons from the first-round L-PCR. Reactions (25 µl, in duplicate or more, using KAPA HiFi HotStart ReadyMix) contained 2.5 µl barcoded primers, dNTPs at 0.6 µM, MgCl_2_ at 2.75 µM, 2% DMSO, and 1.5-3 ng template. For the E amplicons, cycling conditions were the same as those used in the first-round L-PCR, with reduction in the number of cycles to 20. For the G amplicons, conditions were: denaturation at 98°C for 30 s; followed by 30 cycles of 98°C for 10 s, annealing at 70°C for 15 s, and extension 72°C for 6 min; then a terminal extension step at 72°C for 7 min. Conditions for the H amplicon were: denaturation at 95°C for 30 s; 35 cycles of 98°C for 20s, annealing at 74°C for 30 s and extension at 73°C for 5 min; followed by a terminal extension step at 72°C for 10 min. After quantification of amplicons using the same methods as for the first-round, size selection and removal of excess primers was conducted using AMPure PB Beads (PacBio).

Samples were pooled in equimolar amounts (∼23 fmol, in duplicate), calculated using approximations of estimated amplicon lengths and the NEBioCalculator (New England Bio Labs), to generate a pool mass of 0.3-3 μg, with size selection using AMPure PB Beads of the pool, and visualization using the Agilent DNA 12000 kit (Figure 3).

**Figure 3.**
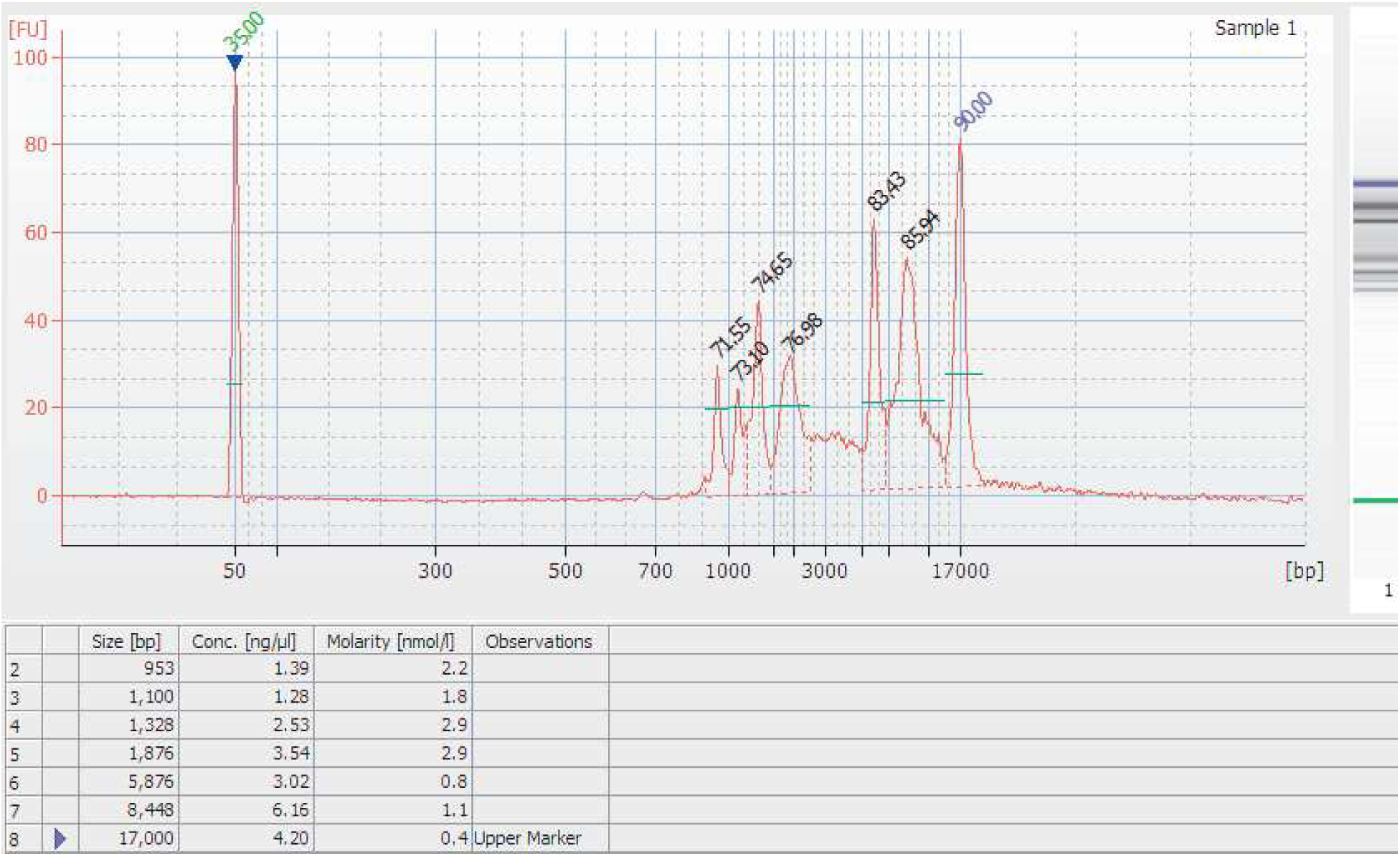
Sample pool as visualized on the Agilent 1200 assay. The ∼5.9 kb peak represents the G amplicons and the ∼8.4 kb peak the E amplicons. The H amplicon are hidden in the region between the G peak and the nonspecific lower peaks.

### Sequencing

SMRTbell library construction using appropriate pool input mass following visualization using an Agilent TapeStation, size selection using AMPure PB Beads, and sequencing with a 30 hour movie time on an 8M cell using the Sequel IIe platform were conducted at The Centre for Applied Genomics (TCAG), the Hospital for Sick Children, Toronto, Canada. The above procedures were also conducted at the Omics and Precision Agriculture Laboratory, Global Institute for Food Security, University of Saskatchewan, where size selection using a Blue Pippin was also conducted. Data demultiplexing was conducted by the bioinformatics teams at these facilities.

### Data Analysis and Sequence Alignment

After filtering CCS reads based on amplicon length, alignment versus reference sequences was conducted using SMRT Link Software version 10.2. Reference sequences (from the National Center for Biotechnology Information (NCBI) or PharmVar) for alignment were deduced from the consensus genotypes previously generated (Supplementary Table S1).

## Results

Amplicons from twenty-one hybrid haplotypes were submitted (for one sample with two types of hybrids, only one was submitted) for sequencing, plus three positive controls. Data from twenty-four hybrid haplotypes were analyzed including positive controls (13 E, 9 G, and two H) (Tables 1-2). For the G amplicons, the alignment versus reference *CYP2D6*13* sequences was 100% (Table 1). The alignment versus reference *CYP2D6*13* sequence for the positive control (NA19785) H amplicon was also 100%. The sample amplified using the H primers that previously had a consensus genotype with a degree of uncertainty aligned best to *CYP2D6*13F* at 99.94%. Technical replicates of this sample all aligned to *CYP2D6*13F* with a percentage of at least 99.5%. The E amplicons aligned to *\*36* and *\*68* hybrid haplotypes as well as to sequences of the various hybrid haplotypes that have the 1847G>A (splice defect) SNV that is the defining SNV for the *\*4* haplotypes (Table 2). Alignment of the *\*68* was 100% and above 99.9% to the *CYP2D6*68* partial sequences EU5300606 and JF307779, respectively, and 99.97 to a sequence derived from the combination of the above partial sequences. The best alignments for the remaining *CYP2D6-2D7* fusion genes were ≥99.7% (to 3 significant figures). The available reference sequences for the *\*4* hybrid haplotypes include EU530605 (*\*4-like*^29^), EU530604 (*\*4N*^29^), and PV00250 (PharmVar *\*4.013*). Alignments against all three of these are presented (Table 2) for the relevant amplicons apart from one, for which alignment against PV00250 was not possible (sample 5).

**Table 1.** Alignment of G amplicons to reference sequences.

| Sample # | Consensus<br>Genotype | Haplotype | Accession<br>number | Filter | Aligned<br>Length | I | D | X | Percent<br>Alignment |
| --- | --- | --- | --- | --- | --- | --- | --- | --- | --- |
| 22 | *13/*4.013 | *13F | EU093102 | 4000 – 6000 | 5002 | 0 | 0 | 0 | 100 |
| 23 | *13/*1 | *13F | EU093102 | 4000 – 6000 | 5002 | 0 | 0 | 0 | 100 |
| 24 | *13/*1 | *13F | EU093102 | 4000 – 6000 | 5002 | 0 | 0 | 0 | 100 |
| 25 | *13+*2/*1 | *13A2 | GQ162807 | 4000 – 6000 | 5039 | 0 | 0 | 0 | 100 |
| 28 | *13+*10/*36 | *13A2 | GQ162807 | 4000 – 6000 | 5039 | 0 | 0 | 0 | 100 |
| 30 | *13/*4.013 | *13F | EU093102 | 4000 – 6000 | 5002 | 0 | 0 | 0 | 100 |
| 32 | *13+*2/*1 | *13A2 | GQ162807 | 4000 – 6000 | 5039 | 0 | 0 | 0 | 100 |
| 33 | *13+*2/*1 | *13A2 | GQ162807 | 4000 – 6000 | 5039 | 0 | 0 | 0 | 100 |
| 35 | *13+*2/*41 | *13A2 | GQ162807 | 4000 – 6000 | 5039 | 0 | 0 | 0 | 100 |
| 37 (NA19785) | *13+*2/*1 | *13A2 | GQ162807 | 4000 – 6000 | 5039 | 0 | 0 | 0 | 100 |

Note: samples 22 and 30 are technical replicates from the same participant. I, D, and X represent the numbers of insertions, deletions, and mismatches, respectively, as compared with the reference sequence.

**Table 2.**
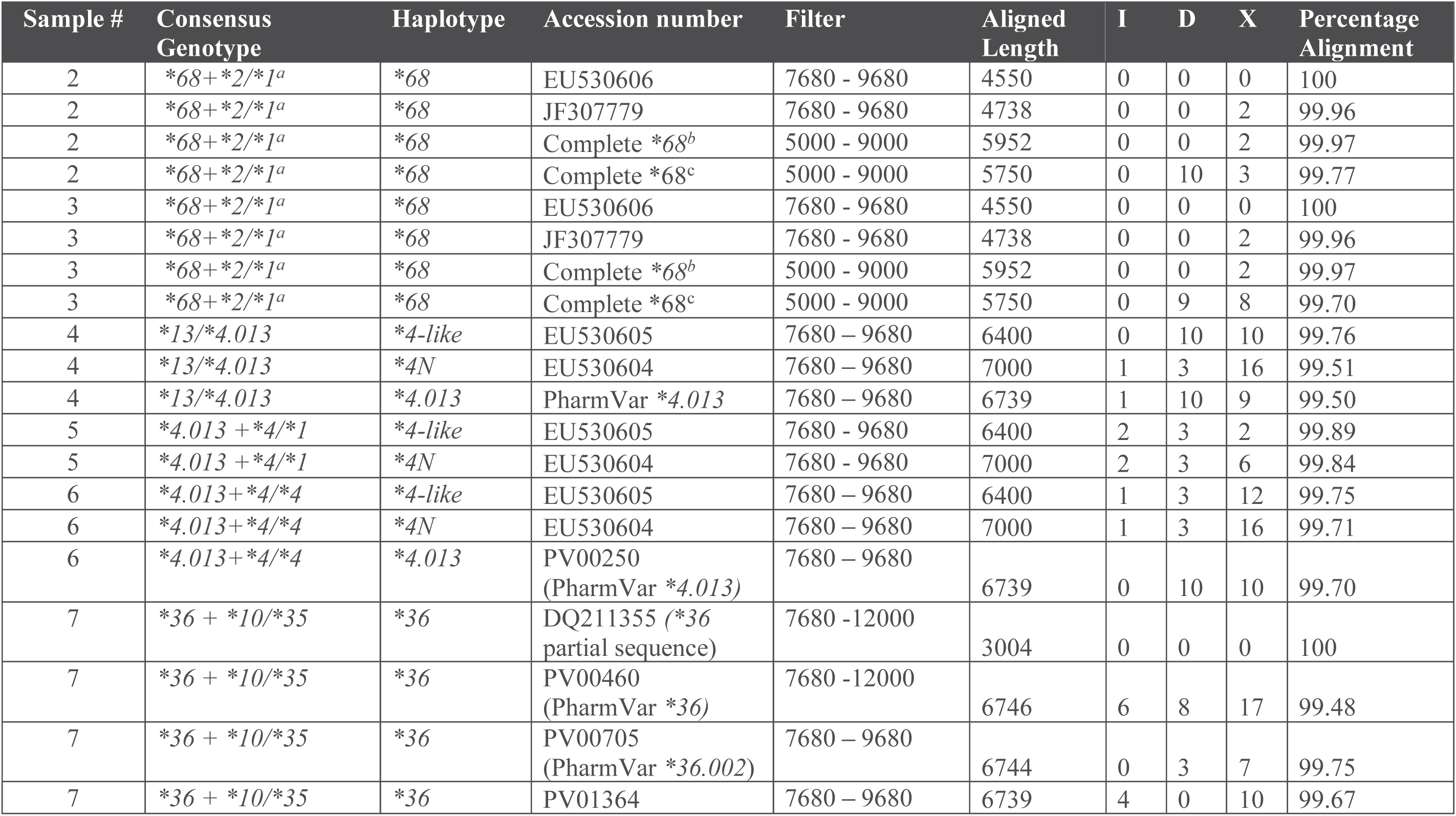
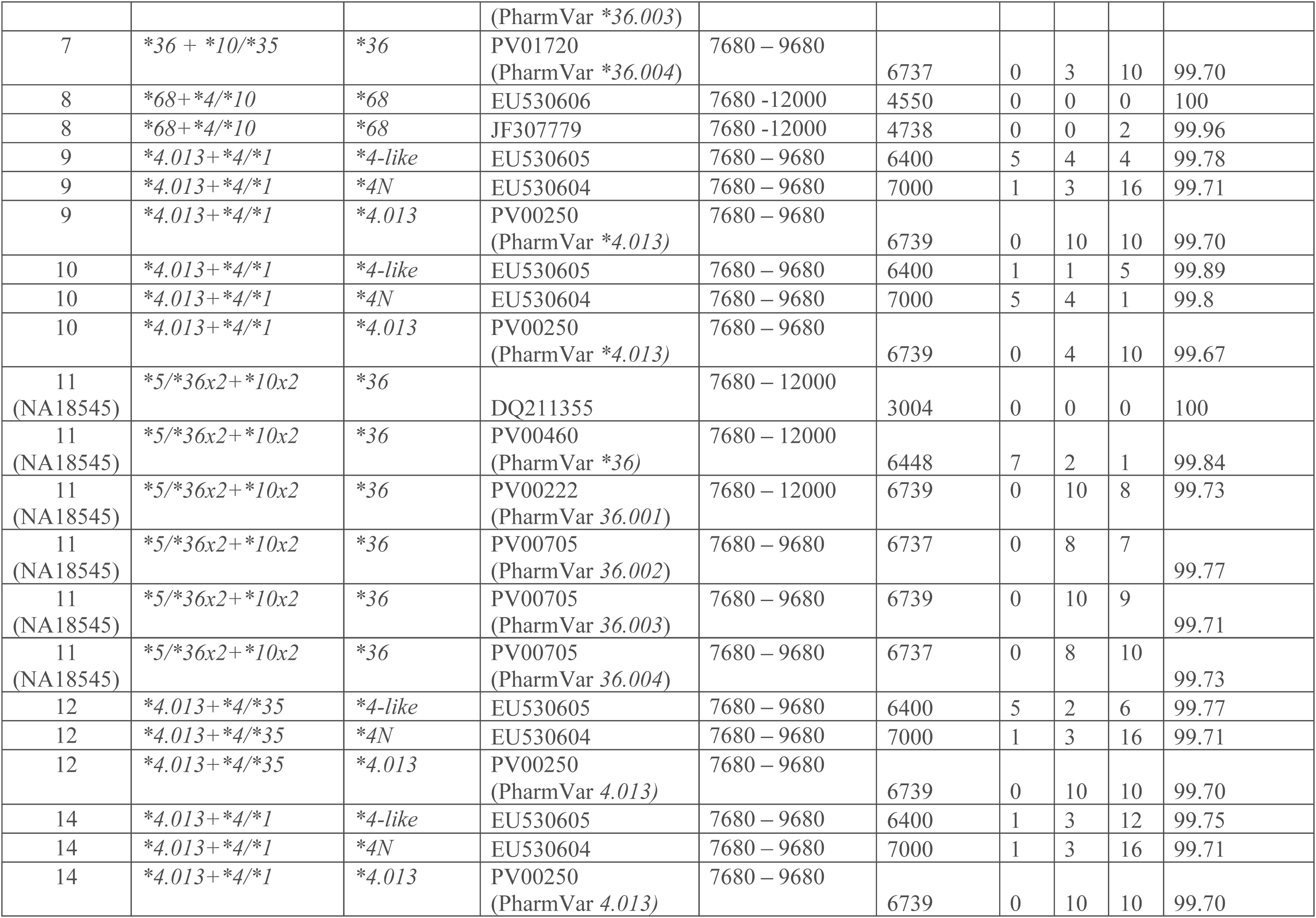
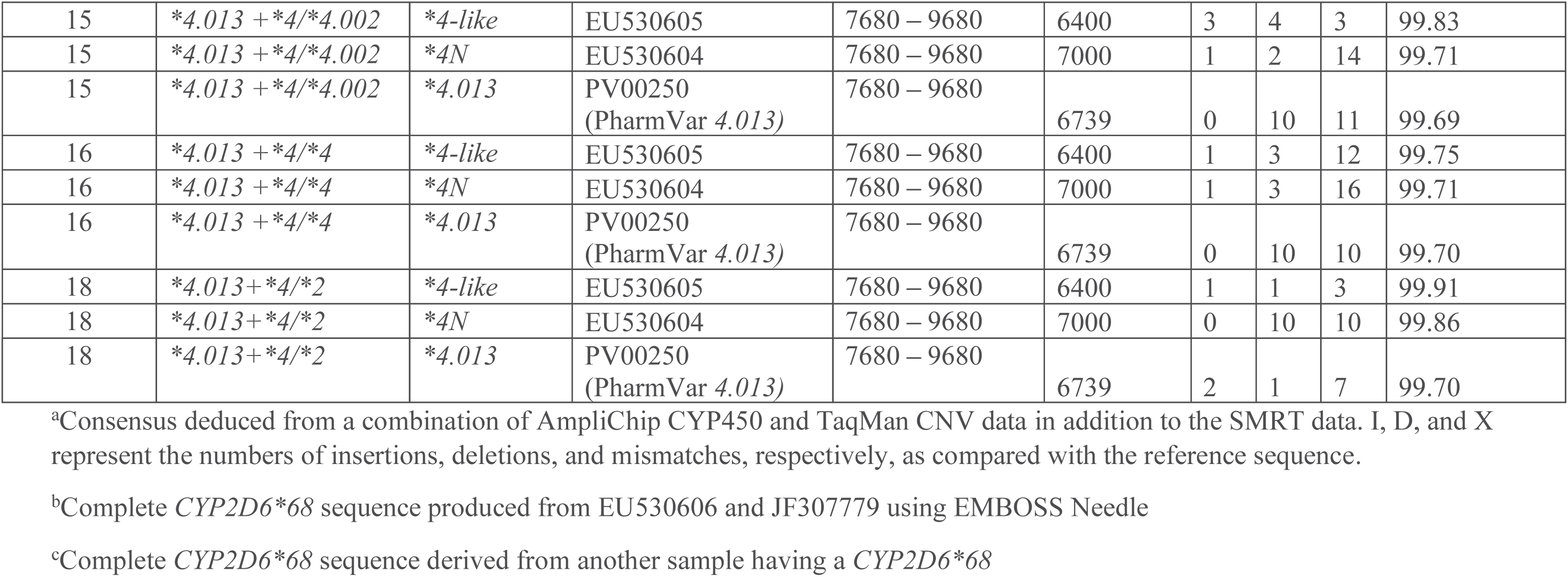
Alignment of E amplicons to reference sequences. For the hybrid haplotypes containing the 1847G>A defining SNV for *4, where possible, alignments to all three reference sequences are provided. For the rest of the samples aligned to more than one reference sequence, unless the percentage alignments were comparable, the best alignment of the longest length is reported. For technical replicates, the best of the replicates is reported.

## Discussion

In summary, we were able to develop and optimize an amplicon-based method of detecting a range of *CYP2D6-2D7* and *CYP2D7-2D6* fusion genes using PacBio barcoded universal primers (BUP) for multiplexing amplicons. This is the most challenging type of *CYP2D6* variant to detect and characterize. Data analysis was highly efficient. Data were cross-validated versus previous data from multiple technologies^43^, and all of the resulting best percentage alignments were ≥99.7%. This method would therefore appear to be more accurate and efficient than any of the other SMRT HiFi methodologies for *CYP2D6* haplotype detection reported to date^40,50,51^ for the detection and characterization of *CYP2D6* fusion genes. In addition, we have since used the forward primer for the E amplicon and the reverse primer for the G amplicon to generate an L-PCR product for non-hybrid *CYP2D6* haplotypes (data not shown). Therefore, the combination of four primer pairs (E forward, E reverse, G forward, G reverse) is sufficient for an amplicon-based method of *CYP2D6* characterization apart from a minority of *CYP2D7-2D6* hybrids for which the H amplicon appears to be required. Moreover, the E forward primer is 1909 base pairs (bp) upstream from the ATG start site, and the G reverse primer 619 bp downstream from the TAG stop site. The amplicon generated therefore covers the upstream (to -1600 bp) and downstream (to 265 bp) regions required for novel haplotype submission to PharmVar.

For G amplicons, the alignment was excellent at 100%. There may be at least two potential reasons for this. Firstly, in the first-round L-PCRs for the Gs, we were able to generate amplicons with minimal non-specificity (Figure 2). Secondly, a *\*13* haplotype was the first type of *CYP2D6* fusion gene to be identified^34^, and hence has been relatively well studied since (with there now being 10 publicly available sequences) in comparison to the other hybrid haplotypes. The percentage alignment for the H positive control was 100%, and the best alignment for the H sample was above 99.9%. Within the E amplicon group, best alignments were at 100% versus partial reference sequences, with the remaining being at ≥99.7%. The greater degree of variability in the alignment statistics for hybrid haplotypes in the *\*4* hybrid haplotypes as compared to other *CYP2D6-2D7* hybrid haplotypes may reflect the fact that (like the various *\*13* hybrid haplotypes^28^) this is a family of hybrid haplotypes containing the 1847G>A SNV. For the samples with a *\*36* haplotype, the alignment statistics for the *\*36* core haplotype and various subhaplotypes were comparable. Sample NA18545 has not previously been sequenced, with the genotype being deduced from CNV testing using L-PCR and quantitative CNV analysis (Gaedigk, personal communication). The alignment to the *\*36* partial sequence was 100% for this sample, and to the PharmVar core *\*36* haplotype 99.84%. As this sample has a *\*36* duplication, it may have more than one *\*36* subhaplotype. Further optimization of the E amplicon procedure as below may contribute to resolving this. Samples with a *\*68* haplotype aligned best to EU530606, with the alignment to the other *\*68* partial reference sequence (JF307779) being 99.96%. Of note, our E amplicons were longer (by ∼1.7 kb) than previously described^29,31^. Initial setting of the extension time in the cycling parameters reflected the shorter predicted length, and while we subsequently set extension to 8 min, this could likely be further optimized to 8.5 min. Accurate amplicon sizing is important not only for L-PCR optimization but also for molar calculations in amplicon pooling. Owing to the presence of some non-specific products at less than the correct length resulting from the E L-PCRs, size selection to remove products less than ∼3 kb was conducted using AMPure PB beads. Whilst the final pool profile indicates the persistence of such products, HiFi sequencing was nonetheless able to produce alignments. This may be at least partly attributable to the relatively small number of multiplexed amplicons, resulting in a high degree of redundancy and read depth.

The main limitation of the work reported herein is the variable percentage alignments, some of which may reflect factors such as PCR optimization achieved to date. Additionally, we recognize that the sample size used here is limited and further work in larger samples is needed for verification of the present methods and results. The alignment previously achieved by Sanger sequencing (after manual curation of the data) was comparable to that achieved by SMRT (e.g., sample 33, alignment 100%; sample 5, alignment 99.75% to *\*4-like*). The present SMRT alignment process was largely automated, and the entire SMRT process reported herein was much more efficient than Sanger sequencing. Further work is in progress, including using more of our own data as reference sequences.

This is the first report of an amplicon-based method of *CYP2D6* SMRT sequencing on a range of fusion genes with haplotypes previously characterized by multiple technologies. Although some versions of *CYP2D6-2D7* fusion genes (e.g., *CYP2D6*61*) were not in our sample set, we are not aware of any reason for our method not working for any *CYP2D6-2D7* fusion gene.

Although CYP2D6 plays a key role in the metabolism of ∼20-25% of clinically used drugs^4,5^, and is included in drug labelling by regulatory bodies (the U.S. Food and Drug Administration, the European Medicines Agency, the Pharmaceuticals and Medical Devices Agency) and prescribing recommendations by consortia (Clinical Pharmacogenetics Implementation Consortium, the Dutch Pharmacogenetics Working Group), identification of *CYP2D6* variants is not yet routine in clinical practice. This is despite the fact that dispensing data indicate that many patients are being prescribed medications for which the identification of *CYP2D6* variants prior to these medications being dispensed could be helpful^55^. One of the reasons for this is the complexity of the locus, and in particular the fact that the fusion genes are challenging to identify and accurately characterize. Another reason is that clinical implementation requires an efficient, high throughput method that requires relatively little personnel time. The HiFi sequencing method reported herein is suitable for high throughput, efficient, with accurate characterization of the full range of *CYP2D6* haplotypes and can be completed in a week or less. Moreover, the method that we have developed could be extended to other loci and might be particularly useful for other complex loci with structural variants, such as *CYP2A6*. In this manner a group of genes may be efficiently characterized in a multiplexed high throughput assay.

## Acknowledgements

We thank the University of Alberta Faculty of Medicine and Dentistry Department of Experimental Oncology and High Content Analysis Core for access to instrumentation and/or services provided, including Michael Wong for his contribution to our Agilent data, and the University of Alberta Department of Biological Sciences Molecular Biology Service Unit (MBSU) for Sanger sequencing (contribution to data in Supplementary Table S1). We thank Andrea Gaedigk for personal communications regarding the details of GeT-RM samples NA19785 and NA18545 used in this report. The following biospecimens donated by [MXL; CHB] were obtained from the NHGRI Sample Repository for Human Genetic Research at the Coriell Institute for Medical Research [NA19785; NA18545]. We thank Matthew Seetin of Pacific Biosciences for bioinformatics training and support and Matthew Ariola (Field Application Specialist) of Pacific Biosciences for technical support. The work described in this paper was funded by: a Canada Foundation for Innovation (CFI), John R. Evans Leaders Fund (JELF) grant (32147—Pharmacogenetic translational biomarker discovery, to KJA), an Alberta Innovates Strategic Research Project (SRP51_PRIME - Pharmacogenomics for the Prevention of Adverse Drug Reactions in mental health; G2018000868 to KJA and Chad Bousman), an Alberta Centennial Addiction and Mental Health Research Chair (to KJA), an Alberta Innovation and Advanced Education Small Equipment Grant (to KJA), the Department of Psychiatry, and the Faculty of Medicine and Dentistry at the University of Alberta. The infrastructure from GP’s lab for the running of the Ion AmpliSeq Pharmacogenomics Panel was supported by a Hotchkiss Brain Institute Dementia Equipment Fund grant (to CB and GP for the Ion) and a Canada Foundation for Innovation John R. Evans Leaders Fund Grant (CFI-JELF) (36624 - Neuromuscular genetics program, to GP). GENDEP was funded by a European Commission Framework 6 grant, LSHB-CT-2003-503428. Roche Molecular Systems supplied the AmpliChip CYP450 Test arrays and some associated support. GlaxoSmithKline and the Medical Research Council (UK) contributed by funding add-on projects in the London centre. This paper is an original work and the views expressed here can only be attributed to the authors, not necessarily reflecting those of the NHS, the NIHR or the UK Department of Health and Social Care.

