## Supplementary Material for "Single molecule long-read real-time amplicon-based sequencing of *CYP2D6*: a proof-of-concept with hybrid haplotypes"

**Supplementary Table S1 Primer Sequences for first-round L-PCR.** Target-specific primer sequences are shown in black; universal forward sequences are shown in blue; universal reverse sequences are shown in green; the 5' amino modifier C6 (5AmMC6) is shown in red.

| Amplicon | Primer | Sequence |
|---|---|---|
| E | Pre alpha F | 5'-5AmMC6GCAGTCGAACATGTAGCTGACTCAGGTCACTCACCCCCAGCGGACTTATCA-3' |
| | Rep 7B rev | 5'-5AmMC6TGGATCACTTGTGCAAGCATCACATCGTAGTACGGTGGGCTCCCTGCGAG-3' |
| G | *16F | 5'-5AmMC6GCAGTCGAACATGTAGCTGACTCAGGTCACCCTGTGTGGGCTTGGGGAGCTTG-3' |
| | *16R5 | 5'-5AmMC6TGGATCACTTGTGCAAGCATCACATCGTAGTGTGGTGAGGTGACGAGGCTGA-3' |
| H | Hyb-F | 5'-5AmMC6GCAGTCGAACATGTAGCTGACTCAGGTCACTCCGACCAGGCCTTTCTACCAC-3' |
| | Hyb-R | 5'-5AmMC6TGGATCACTTGTGCAAGCATCACATCGTAGCGACTGAGCCCTGGGAGGTAGGTAG-3' |
